# Effect of soilless media on small-scale propagation of *Pinguicula gigantea*

**DOI:** 10.1101/2023.01.18.523674

**Authors:** Kevin Tarner, Hakon Jones, Mile Ingwers, Heather Gaya, Mason McNair

## Abstract

Because of their large size, colorful flowers, and insectivorous habit, butterworts (genus *Pinguicula)* are desirable ornamental plants among hobbyists and botanical conservatories. Propagation via leaves is a popular propagation method for the genus, especially for tropical species. *P. gigantea*, a species found approximately 700m above sea level in Oaxaca, Mexico, was selected to evaluate its preference for various blends of soilless media. This study found that number of leaves produced by plantlets is significantly impacted by soil type. However, there was no significant difference in biomass, plantlet diameter or average number of plantlets produced between soil treatments. These results suggest that soils with high nitrogen content may promote increased leaf number, but do not significantly affect plant biomass.

## INTRODUCTION

*Pinguicula* are cultivated for their ornamental value, insectivorous nature, and for conservational purposes. Of the 105 recognized *Pinguicula* (Lentibulariaceae) species, 15 are critically endangered, 4 are endangered, 31 are vulnerable, 3 are near threatened, and there are insufficient data to evaluate another 19 species (Cross et al., 2020). *Pinguicula gigantea* is a member of the largest section of the genus (sect. Temnoceras) and is found at approximately 700m above sea level in Oaxaca, Mexico (Fleischmann & Roccia, 2018). This species is unique among the genus in that it prefers airy, well-draining soil, a stark contrast to the wet habitats preferred by related species (Lampard et al., 2016). *P. gigantea* reproduces by seed and by forming clonal plantlets on leaves that have detached from a mother plant. The species’ large size in comparison to other species and showy purple or pink flowers make it desirable among hobbyists and botanical conservatories. Poaching is one of the largest threats facing carnivorous plants today (Lowther et al., 2002). Inexpensive, low-tech, and easy clonal propagation of *Pinguicula* may help reduce demand for poached specimens. Past research has focused on mass propagation of *Pinguicula* species via in vitro propagation (Clapa et al., 2010; Gonçalves et al., 2008). This type of propagation is rarely accessible to smaller nurseries, hobbyists, and conservationists; groups which play prominent roles in market demand and conservation efforts for important carnivorous plants like *Pinguicula*. The hobby community is full of successful and diverse anecdotal evidence surrounding propagation of *Pinguicula*, including non-peer-reviewed publications (Brittnacher, 2019; D’Amato, 1998; *Growing Guides* | *ICPS*, 2022). However, few peer-reviewed studies on the effect of soil or plant nutrient content or fertilizer on carnivorous plants exists, especially for the genus Pinguicula (Adamec, 2002; Adlassnig et al., 2012; Capó-Bauçà et al., 2020; Dixon et al., 1980; Ellison, 2006; Ellison & Gotelli, 2002; Gao et al., 2015; He & Zain, 2012).

This paper focuses on the propagation of a single clone of *P. gigantea* tested in three easily reproducible substrates in a common greenhouse environment at the University of Georgia (UGA). The goals of this study were to evaluate the role of substrate in *Pinguicula* propagation success and post-propagation plantlet growth. It was hypothesized that tropical *Pinguicula* might exhibit plasticity in acceptable growing media preference, regarding properties like drainage, particle size, pH, and amendment with chemical fertilizers.

## MATERIALS AND METHODS

*Pinguicula gigantea* leaves were collected from mature, flowering-sized plants in the UGA Plant Biology Greenhouse Teaching Collection (Figure 1A). All *P. gigantea* in this collection consist of a single clone originally obtained from John Brittnacher in April, 2014. This material was particularly suitable for this experiment because of the large quantity of leaves available as well as the ability to select for uniformity of leaf size. Substrates were mixed prior to the beginning of the experiment using established protocols within the UGA Plant Biology Greenhouses (Table 1).

**Table 1.**
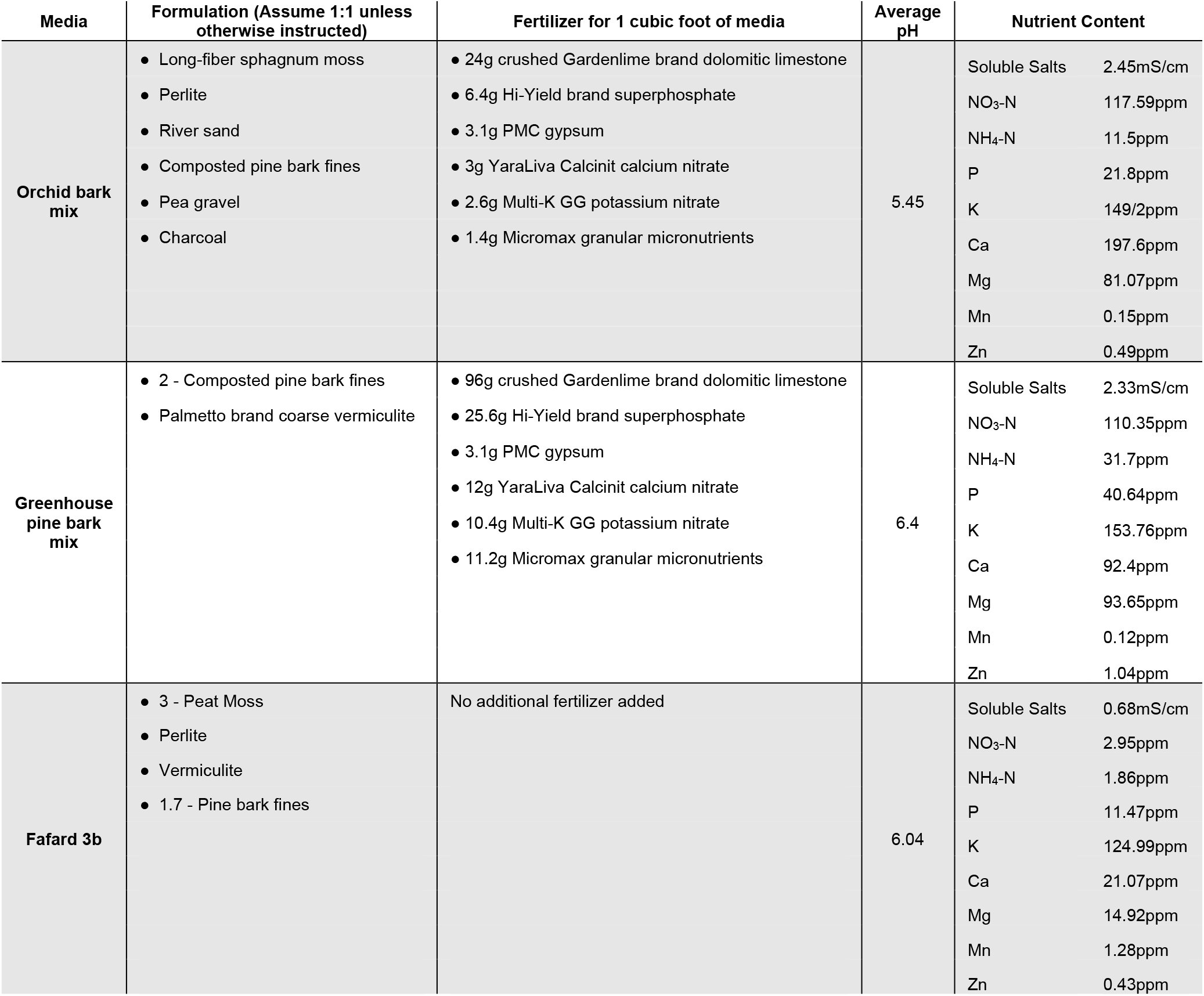
Soilless media formulations. Media formulations assume equal parts unless otherwise stated. Units for fertilizer are in grams (g).

**Figure 1.**
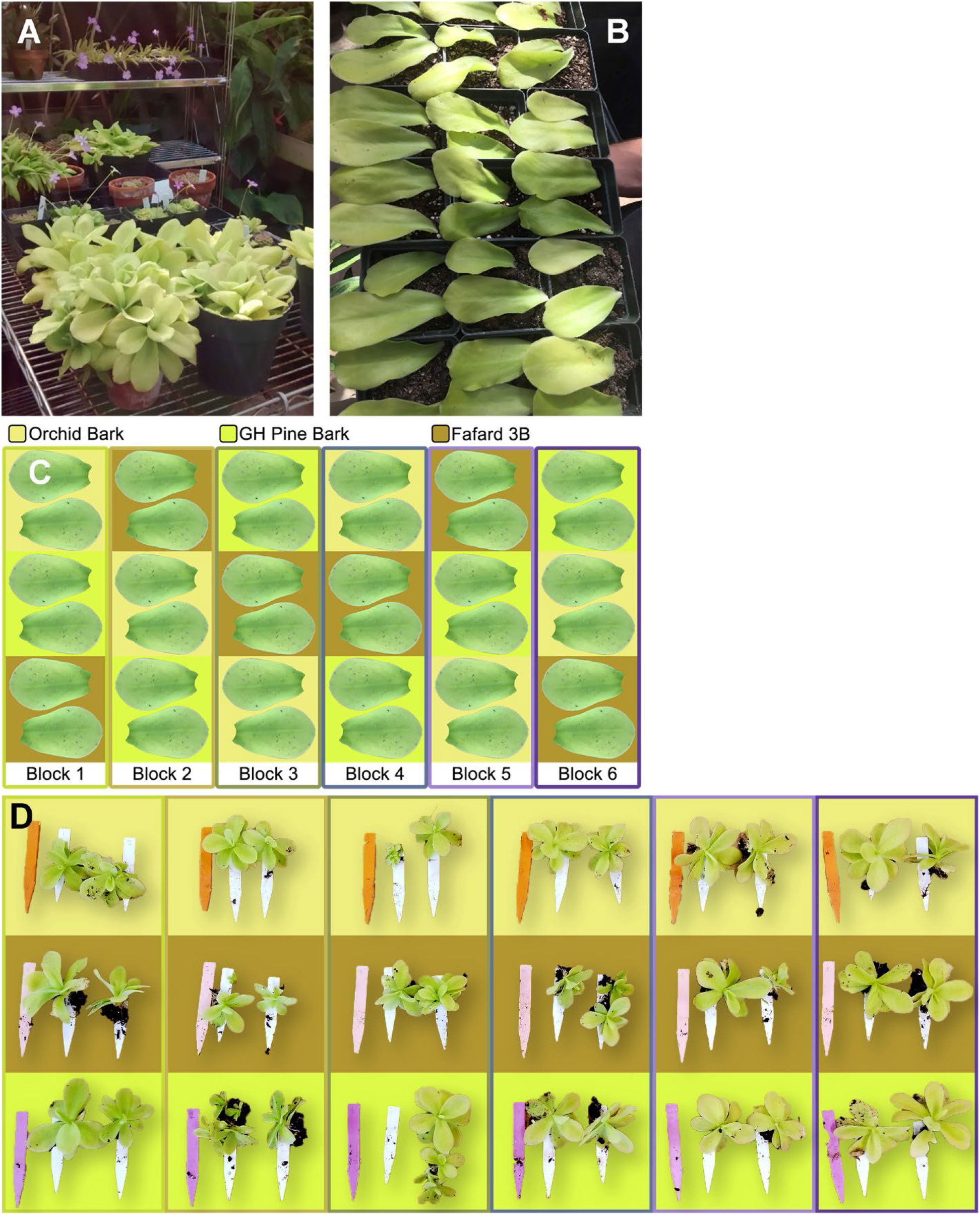
A. Stock plants of *P. gigantea* used in this experiment. All plants are one clone propagated from leaf cuttings since 2014. B.36 leaves were selected for approximate size uniformity, grouped into three treatments consisting of six randomized blocks with 2 replicates per block and laid on the surface of the substrate. C.Experimental Design showing treatments, blocks, and replicates. D.Propagated plantlets organized by media type (rows) and by block (columns). Colored tags are 12.7cm long. White tags are plantlets from each replicate.

Leaves were pulled gently from multiple specimens of this single clone, selected within each block for approximate uniformity of size, and laid on the surface of different soilless medias. A total of 36 leaves were harvested from stock clones and grouped into 3 treatments (Fafard3b, OrchidBark, GHPineBark), each consisting of six randomized blocks (4-inch pots) with two replicates per block. Treatments were labeled using numbered tags and placed into a 10×20 plastic flat (Figure 1B). Each leaf was weighed immediately after being harvested from the stock plant. Treatments were placed into the greenhouse at 11AM October 25th, 2017. Harvest occurred when it was determined that all the original leaf pullings had completely senesced (2pm, April 14th, 2018), a total duration of 192 days or approximately 27 weeks. At harvest, the number of plantlets, leaf number per plantlet, and plantlet diameter were measured (Table 2). All media was removed from the roots and every individual was bulked by replicate, labeled, and dried to a constant mass before final weight measurement was taken.

**Table 2.** Collected Data. All data was collected manually and averaged (when applicable) prior to statistical analysis.

| Plantlet Number | Average Leaf Number | Average Plantlet Diameter (millimeters) | Initial Weight (grams) | Dry Weight (grams) | Treatment | Block |
| --- | --- | --- | --- | --- | --- | --- |
| 1 | 11 | 80 | 2.86 | 0.183 | OrchidBark | 1 |
| 2 | 9 | 72 | 3.88 | 0.2334 | OrchidBark | 1 |
| 1 | 12 | 82 | 4.48 | 0.2375 | OrchidBark | 2 |
| 1 | 9 | 74 | 1.63 | 0.1233 | OrchidBark | 2 |
| 7 | 8.571428571 | 17.28571429 | 3.73 | 0.0408 | OrchidBark | 3 |
| 1 | 10 | 70 | 2.96 | 0.1232 | OrchidBark | 3 |
| 1 | 12 | 88 | 4.68 | 0.2485 | OrchidBark | 4 |
| 1 | 11 | 82 | 1.57 | 0.1843 | OrchidBark | 4 |
| 1 | 12 | 95 | 3.58 | 0.2176 | OrchidBark | 5 |
| 1 | 12 | 107 | 2.47 | 0.315 | OrchidBark | 5 |
| 1 | 11 | 106 | 3.34 | 0.3061 | OrchidBark | 6 |
| 1 | 12 | 83 | 2.33 | 0.1722 | OrchidBark | 6 |
| 1 | 11 | 123 | 2.86 | 0.3515 | Fafard313 | 1 |
| 1 | 9 | 95 | 2.07 | 0.1711 | Fafard313 | 1 |
| 2 | 8.5 | 44.5 | 3.05 | 0.1187 | Fafard313 | 2 |
| 1 | 9 | 61 | 1.73 | 0.0646 | Fafard313 | 2 |
| 1 | 11 | 85 | 1.80 | 0.159 | Fafard313 | 3 |
| 2 | 10.5 | 66.5 | 4.19 | 0.1969 | Fafard313 | 3 |
| 1 | 9 | 80 | 1.19 | 0.1055 | Fafard313 | 4 |
| 2 | 8 | 78 | 2.45 | 0.1899 | Fafard313 | 4 |
| 1 | 11 | 109 | 2.16 | 0.339 | Fafard313 | 5 |
| 3 | 8.666666667 | 35 | 4.19 | 0.0949 | Fafard313 | 5 |
| 1 | 11 | 100 | 3.08 | 0.2658 | Fafard313 | 6 |
| 1 | 11 | 92 | 4.44 | 0.3254 | Fafard313 | 6 |
| 1 | 12 | 117 | 4.00 | 0.3397 | GHPineBark | 1 |
| 1 | 12 | 78 | 4.47 | 0.1164 | GHPineBark | 1 |
| 4 | 9.75 | 53.75 | 3.51 | 0.2148 | GHPineBark | 2 |
| 1 | 12 | 85 | 3.50 | 0.1901 | GHPineBark | 2 |
| 2 | 10.5 | 67.5 | 2.04 | 0.1512 | GHPineBark | 3 |
| 1 | 12 | 91 | 2.28 | 0.3003 | GHPineBark | 3 |
| 1 | 11 | 99 | 3.66 | 0.2486 | GHPineBark | 4 |
| 1 | 11 | 96 | 3.74 | 0.211 | GHPineBark | 4 |
| 1 | 10 | 88 | 2.17 | 0.2259 | GHPineBark | 5 |
| 1 | 11 | 83 | 3.33 | 0.1789 | GHPineBark | 5 |
| 1 | 12 | 97 | 4.19 | 0.212 | GHPineBark | 6 |
| 1 | 11 | 110 | 2.73 | 0.3515 | GHPineBark | 6 |

Environmental data from the greenhouse room containing this experiment, consisting of temperature and relative humidity, was logged for the duration of the experiment using a Lascar Electronics model EL-USB-2-LCD datalogger. The average temperature was 18.56°C, fluctuating between 9°C and 27°C throughout the experiment. The average relative humidity was 62.22%, fluctuating between 22.5% and 95%. The average dewpoint was 10.84°C, fluctuating between -4.2°C and 22.5°C.

Results were analyzed using an ANCOVA run in RStudio (R Core Team, 2019) to determine if soil treatment had a significant effect (p<0.05) on number of plantlets, plantlet dry weight, plantlet diameter, or number of leaves per plantlet. All ANCOVA were tested for the effect of initial weight of leaf pullings.

## RESULTS

Substrate did not significantly affect the number of plantlets produced (p=0.872), plantlet diameter (p=0.559), or dry weight (p=0.612) of the samples (Figure 2). However, the average number of leaves per plantlet was significantly affected by media (p = 0.016). Results did not change when all models were run with an adjustment for initial leaf weight (p = 0.629, p = 0.6738, p = 0.638, and p = 0.019 respectively).

**Figure 2.**
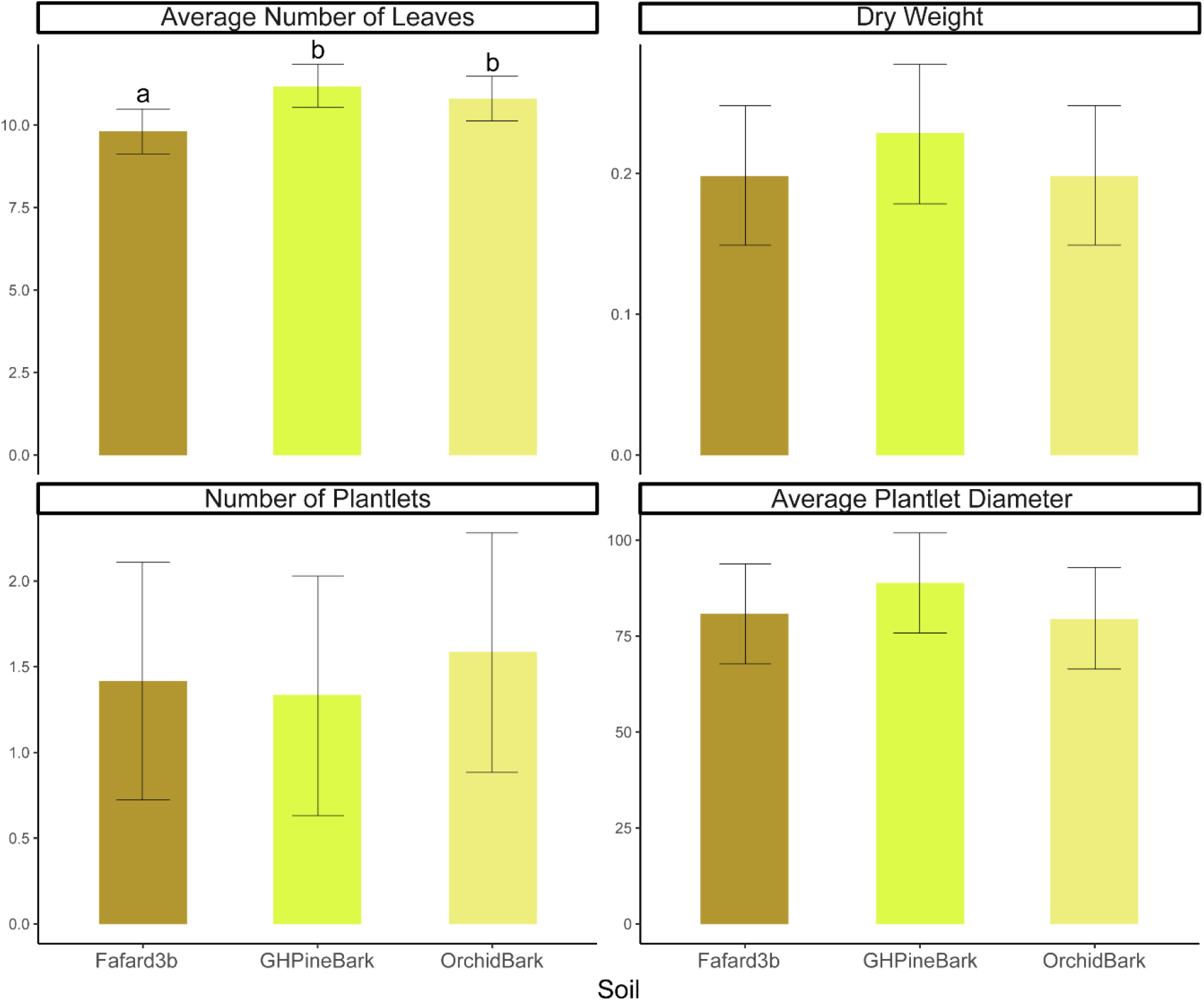
Comparison of the average number of leaves, dry weight, number of plantlets, and plantlet diameter for 36 *P. gigantea* grown under three soil conditions. Specimens grown in Greenhouse Pine Bark and Orchard Bark produced significantly (p < .05) more leaves per plantlet than those in Fafard 3b (A). There were no significant differences found in dry weight (B), number of plantlets (C), or average plantlet diameter (D) between plants grown in the three soil types.

The Greenhouse pine bark mix produced significantly more leaves per plantlet than the Fafard 3b substrate (one-sided t-test, df = 19.4, p = 0.002) and Orchid bark mix produced significantly more leaves per plantlet than Fafard 3b (one-sided t-test, df = 21.8, p = 0.034, Figure 3). There was no significant difference in the average number of leaves per plantlet between Greenhouse pine bark and Orchid bark mix (one-sided t-test, df = 18.3, p = .199).

**Figure 3.**
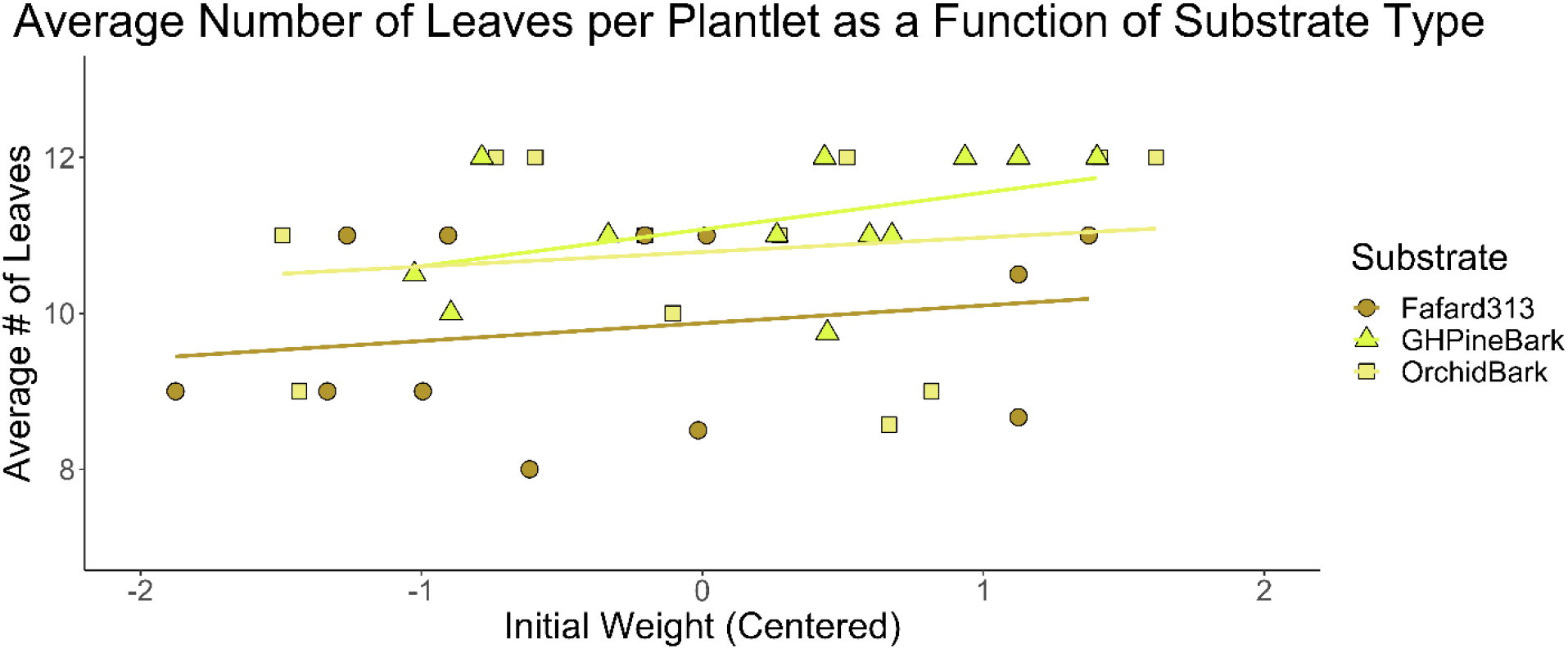
The average number of leaves per plantlet, based on initial weight and grouped by substrate type. The average number of leaves was significantly lower for the Fafard313 mix relative to the other treatments, even after accounting for initial pulling weight.

## DISCUSSION & CONCLUSION

*Pinguicula gigantea* stock plants produce masses of fallen leaves and fall randomly into pots below with OrchidBark, Fafard 3B, or Greenhouse pine bark mixes in the UGA Plant Biology Teaching collection. It was hypothesized that *P. gigantea* might exhibit significant plasticity with regards to growth and propagation in multiple substrates. Notably, all substrates used in this experiment are circumneutral in pH (between 5.5 and 7.2), supplemented with a variety of chemical fertilizers (all proving non-phytotoxic in the long term), variable in particle size and drainage, and intended for the growth of non-carnivorous plant taxa. A nutrient devoid soilless media was not used as a control despite the carnivorous nature of *P. gigantea*. It is a common misconception that carnivorous plants can survive without any soil nutrition. No outright deaths of leaves or plantlets occurred at any point during this study, supporting our hypothesis about *Pinguicula* plasticity in cultivation; however further research is needed to statistically support this hypothesis. It is well documented that carnivorous plants grow in poor nutrient soils but peer-reviewed studies investigating the use of soilless media and inorganic fertilization are few (Adamec, 2008; Zamora et al., 1997). However, anecdotal evidence is abundant in both printed and online literature (Brittnacher, 2019; D’Amato, 1998; *Growing Guides* | *ICPS*, 2022).

Both Greenhouse pine bark mix and OrchidBark (formulations in Table 1) provided significant increases in average number of leaves per plantlet, but not in final biomass, average plantlet diameter, or number of plantlets per pulling (Figure 2). This is likely explained by the large amount of fertilizer added to these substrates, providing initial nutrients to plantlets as they senesce from the mother leaf pulling and root into the substrate. Both the Greenhouse and OrchidBark mixes contained high levels of soluble salts, NO_3_-N and NH_4_-N relative to the Fafard 3b mixture (Table 1). While increased substrate nitrogen appears to have a positive effect on the number of leaves per plantlet, additional substrate nutrients may be necessary to increase final biomass or average plantlet diameter. Future research focusing on mineral uptake of individual plants could allow for a better understanding of the relationship between substrate nutrients and plant growth.

The findings herein may also translate to other *Pinguicula* species, potentially allowing for increased propagation, cultivation, and conservation. Future studies should consider examining the effects of common fertilizers on growth of *P. gigantea* and replicating this study to include larger sample sizes and additional taxa.

## ACKNOWLEDGEMENTS

The authors would like to thank John Brittnacher for providing the clone of *Pinguicula gigantea* utilized in these experiments. We would also like to thank Dr. James Leebens-Mack for their insightful review and commentary on this manuscript prior to submission.

## Notes

### Competing Interest Statement

The authors have declared no competing interest.

